# Obesogenic diet and targeted deletion of potassium channel K_v_1.3 have differing effects on voluntary exercise in mice

**DOI:** 10.1101/650978

**Authors:** Brandon M. Chelette, Abigail Thomas, Debra Ann Fadool

## Abstract

Voluntary exercise is frequently employed as an intervention for obesity. The voltage-gated potassium K_v_1.3 is also receiving attention as a therapeutic target for obesity, in addition to potential therapeutic capabilities for neuroinflammatory diseases. To investigate combinatorial effects of these two therapies, we have compared the metabolic status and voluntary exercise behavior of both wildtype mice and a transgenic line of mice that are genetic knockouts for K_v_1.3 when provided with a running wheel and maintained on diets of differing fat content and caloric density. We tracked metabolic parameters and wheel running behavior while maintaining the mice on their assigned treatment for 6 months. Wildtype mice maintained on the fatty diet gain a significant amount of bodyweight and adipose tissue and display significantly impaired glucose tolerance, though all these effects were partially reduced with provision of a running wheel. Similarly to previous studies, the K_v_1.3-null mice were resistant to obesity, increased adiposity, and impaired glucose tolerance. Both wildtype and Kv1.3-null mice maintained on the fatty diet displayed increased wheel running activity compared to CF-fed mice which was caused primarily by a significant increase in amount of time spent running as opposed to an increase in running velocity. Interestingly, the patterns of running behavior differ between wildtype and K_v_1.3-null mice, especially in how their resting periods are distributed through the dark phase. These studies indicate that voluntary exercise combats metabolic maladies and running behavior is modified by both consumption of an obesogenic diet and deletion of the K_v_1.3 channel.

**NEW and NOTEWORTHY:**

- K_v_1.3-null mice exhibit different running and resting patterns compared to wildtype mice
- Mice maintained on an obesogenic diet (32% kcal from fat) exhibit increased running distance and increased time spent running compared to mice fed normal rodent chow.

## INTRODUCTION

The prevalence of obesity in the human population is increasing at an alarming rate. In the United States, the percentage of obese adults has increased from 15% in 1990 to over 35% in 2012 and obesity is now commonly described as an epidemic {1}. Obesity has been correlated with increased incidence of many diseases as well as increased mortality {2}. Financial impact estimates vary, but the cost of obesity is reported in the hundreds of billions of dollars in the United States {2}. These troubling trends have necessitated an increased research effort to uncover prevention and treatment strategies for obesity.

Exercise has long been prescribed to treat obesity, but the benefits of engaging in voluntary exercise extend beyond attempts to lose weight {3}. Studies have indicated that participation in voluntary exercise suppresses inflammation in the central nervous system, slows the cognitive decline and reduction of hippocampal volume associated with aging, modulates the microbiome, and combats insulin resistance {4,5,6}. Conversely, inactivity has been identified as a risk factor for the development of neurodegenerative diseases such as Alzheimer’s disease and dementia {7}. In light of the obesity epidemic, the more recent discovery of the additional health benefits of exercise, and the detriments of a sedentary lifestyle, voluntary exercise is receiving renewed research attention {8,9}.

Another promising therapeutic target is the voltage-gated potassium channel, K_v_1.3. One of the largest families of ion channels, potassium channels were historically characterized to function as dampeners of excitability through timing of the interspike interval and shaping of the action potential {10}. We now know there are multifarious roles of potassium channels that include a number of non-conductive functions such as their ability to regulate or detect energy substrates, to be key players in immune responses, and to modulate sensory perception {11,12,13}. K_v_1.3 is a mammalian homolog of the *Shaker* subfamily that has a select distribution within the CNS (the olfactory bulb, the pyriform cortex, and the dentate gyrus of the hippocampus), the peripheral organs (adipocytes, kidney), and the immune system (T-cells) {14,15,16,17,18,19,20}. Efforts by our laboratory and others have revealed that K_v_1.3-null mice exhibit several interesting metabolic phenotypes. Multiple studies have shown that K_v_1.3-null mice are thin compared to wildtype mice, have a resistance to diet-induced obesity (DIO), and exhibit a slightly increased basal metabolic rate along with increased locomotor activity {21,22,23}. Further, blockade of K_v_1.3 increases glucose uptake by increasing the trafficking of glucose transporter type 4 (GLUT4) to the membrane of skeletal muscle cells and adipocytes {18}. Gene deletion or pharmacological blockade of K_v_1.3 has been shown to reduce neuroinflammation and has been used to treat a range of animal model versions of autoimmune diseases including multiple sclerosis, psoriasis, and type-1 diabetes {24}. The treatments have been effective and researchers have been successful in increasing the potency, selectivity, and targeting of K_v_1.3 blockers (peptides isolated from scorpion and sea anemone toxins) {25}. Due to this unique combination of physiological effects driven by K_v_1.3, the channel has become an important and promising therapeutic target for neuroinflammatory diseases, metabolic disorders, and neurodegeneration {26,27}.

The most promising therapeutic directions for both voluntary exercise and K_v_1.3-targeted treatments are converging on a subset of human diseases for which both treatments might be effective. This is especially true for diseases that have both a metabolic and autoimmune component. Progress is being made rapidly in both areas of research, but little is known concerning the potential combinatorial effects of these two therapeutic interventions. Further, although the biophysical properties of K_v_1.3 in many cells are well characterized and the channel has become an important therapeutic target as outlined above, less is known concerning how loss of K_v_1.3 may affect locomotor activity patterns or if voluntary running is enhanced in the absence of the channel.

Our investigation is the first to characterize the running phenotype of mice where we measured the interactions of six-month access to voluntary exercise, diet modification, and genetic loss of the K_v_1.3 channel. We found that both wildtype and K_v_1.3-null mice challenged with a moderately high-fat diet participated in wheel running more often and ran farther than control-fed mice. We also demonstrated that participation in voluntary exercise, as expected, partially protected wildtype mice from increased adiposity and glucose insensitivity associated with the MHF diet when compared to MHF-fed sedentary mice. Finally, K_v_1.3-null mice ran the same distance and velocity as wildtype mice when maintained on the same diet, but their patterns of resting and running differed across the both the light and dark phases.

## MATERIALS AND METHODS

### Animal Care

All mice were maintained in the Florida State University (FSU) animal vivarium in accordance with the institutional requirements established by the FSU Animal Care and Use Committee (ACUC) and all experimental procedures were performed according to the approved FSU ACUC protocol #1733. Our lab and others had previously demonstrated that female mice did not gain adiposity (lacked diet-induced obesity, DIO), therefore only male mice were used in this current investigation to explore the intersection of diet, voluntary exercise, and presence/absence of K_v_1.3 channel {23}. Mice were weaned to large, breeding-style cages (18 x 9.25 x 6.25 cm) at postnatal day (PND) 21-25 to accommodate the running wheel equipment (see **Voluntary Exercise Behavior** section for a description of the wheel equipment). The mice were maintained on either a normal rodent chow diet (Purina 5001 Rodent Chow; 13.5% kcal from fat) or a moderately high-fat (MHF), condensed milk diet (Research Diets, Cat #D12266B, 31.8% kcal from fat). The mice were singly-housed in conventional cages upon weaning, provided *ad libitum* access to food and water, maintained on a 12h/12h light/dark cycle, and provided only their assigned diet for the duration of the experimental period (~160 days).

### Animal Lines

To investigate the role of the ion channel K_v_1.3 on exercise behavior, a transgenic line of mice that were global genetic knockouts for K_v_1.3 (K_v_1.3-null) were used. These mice were on a C57BL/6J background and were generated via deletion of a large promoter region as well as the N-terminal third of the coding sequence for K_v_1.3 {22,28}. The mice were a generous gift of Drs. Leonard Kaczmarek and Richard Flavell (Yale University, New Haven, CT) and have now been deposited at Jackson Laboratories (Bar Harbor, ME; B6;129S1-Kcna3tm1Lys/J, stock number 027392). The K_v_1.3-null mice (KO) were compared to mice with a wildtype (WT) version of the channel. The wildtype cohort consisted of two lines of mice. The first were C57BL/6J mice from Jackson Laboratories (stock number 000664). The second were *M72-IRES-tau-Lac* mice (gene *oflr160*) on a C57BL/6J background. The only difference between the C57BL/6J and the *M72-IRES-tau-LacZ* mice was an addition of a genetic reporter in the latter, which allows for the visualization of a subset of olfactory sensory neurons {29,30,31}. The *M72-IRES-tau-LacZ* mice were generously provided by Dr. Peter Mombaerts (Max-Planck-Gesellschaft, Munich, Germany) and were used here with the intention of applications for future experiments to investigate the anatomy of the olfactory system following voluntary exercise and pair feeding (not performed here) {32} Several investigators have shown that voluntary exercise is highly variable and certain transgenic lines run abnormal amounts when given access to a wheel {33}. However, there were no differences between the running behavior (or any other metrics) of the C57BL/6J vs the *M72-IRES-tau-LacZ* mice, so these two groups were pooled and referred to as wildtype (WT) throughout this manuscript.

### Bodyweight and Adiposity

To compare normal weight gain to the increased weight gain and deposition of adipose tissue caused by maintenance on the MHF diet, the bodyweight of each mouse was recorded weekly. Upon termination, fat pads were excised and weighed by an investigator that was blind to the genotype and treatment group of the mouse. Endometrial, retroperitoneal, subcutaneous (a subsample of the right side), and mesenteric adipose tissues were collected and combined. Brown fat was not sampled.

### Glucose Tolerance

Mice were fasted for 12 hours through their dark phase (0800h – 2000h) prior to administration of an intraperitoneal glucose tolerance test (IPGTT). Mice were injected with a volume of 25% glucose solution equivalent to 1 g of glucose per kg of bodyweight (University of Virginia Vivarium Protocols, Susanna R. Keller). A small incision was made on the tail and blood samples were collected with an Ascensia CONTOUR™ Blood Glucose Monitoring System (Bayer Healthcare, Whippany, NJ) paired with CONTOUR™ Blood Glucose Test Strips (Bayer Healthcare) to determine blood glucose levels at baseline (prior to injection) and at set timepoints 10, 20, 30, 60, 90, and 120 minutes following the injection.

### Voluntary Exercise Behavior

Mice belonging to the voluntary exercise treatment groups were provided *ad libitum* access to a running wheel in their cage. Following a 2-day acclimation period to their larger, individual cage, a Vertical Wireless Running Wheel (Med Associates, Inc., St Albans, VT) was affixed in the cage. The wheel sensor was placed atop the wire lid and plastic manifolds extended into the cage that held the running wheel above the bedding and allowed for rotation. A clean wheel was provided weekly and the wheels were lubricated with non-caloric, tasteless, and odorless silicone oil as necessary to minimize potentially stressful squeaking of the rotating wheel. The wheel sensor recorded the number of rotations and transmitted the data to a central hub every 30 seconds (s). The hub was in turn connected to a computer equipped with the Wheel Manager software program (Med Associates, Inc.), which archived the data and allowed for data exportation upon conclusion of the running period. The running data were exported to Microsoft Excel via the Wheel Analysis software program (Med Associates, Inc.). A 1-minute bin was the smallest bin size that the software allowed to be exported.

A majority of the running behavior data reported in this manuscript are a 28-day subsample for each mouse instead of the entire experimental period of 4 months. We have concluded that restricting analysis to a 28 day period of only the dark phase is justifiable because 1) it allowed for the exclusion of the 3-day acclimation period during which mice ran very little and/or inconsistently and 2) it allowed for the exclusion of outliers in running behavior that were obviously due to a technical error (trapping of bedding or low battery) as opposed to actual behavioral changes of a mouse.

Mean running distance was calculated as the daily average of kilometers run per day across the 28-day sample for each mouse. Active time was calculated as the number of 1-minute bins during which the mouse was participating in wheel running. Running velocity was reported as the mean wheel rotations per minute for every 1-minute bin during which the mice was active. Running bursts were defined as consecutive 1-minute bins during which the mouse was active while rests were defined as consecutive 1-minute bins during which the mouse was inactive. Latency to run is defined as the mean amount of time after dark onset until a mouse engaged in wheel running. Latency to stop is defined as the mean amount of time after light onset until a mouse stopped engaging in wheel running.

### Statistical Analysis

Final body weight, weight of adipose tissue, integrated area-under-the-curve analysis of glucose tolerance, and all the calculated running behaviors (distance, velocity, bursts, etc.) were compared using an analysis of variance (ANOVA). All statistical tests used α = 0.05 as the minimum confidence interval for significance and all reported values are mean ± standard deviation (s.d.). All post-hoc analyses utilized the Tukey’s multiple comparison test and significantly different mean-wise comparisons were indicated by different letters in the figures. Data organization and analysis was performed with Microsoft Excel and The R Project statistical computation system {34}. Double-plotted actograms were generated using Fiji and the associated plugin ActogramJ {35,36}. Graphs were designed and generated using Origin Student 2018b software (version b9.5.5.409 OriginLab Corporation, Northhampton, MA).

Statistical tests were performed with Graphpad Prism (version 7.04, Graphpad Software, La Jolla, CA).

## RESULTS

### Wheel running offers partial protection from the effects of MHF-diet

WT mice fed the CF diet weighed the same regardless of whether they had access to a running wheel (WT-CF-SED 24-week bodyweight = 27.8 g ± 2.1; WT-CF-RW = 28.0 ± 1.4). Mice fed the MHF diet without access to a running wheel gained significantly more bodyweight than all other treatment groups (WT-MHF-SED = 39.2 g ± 4.4; F(7,77) = 42.2; p < 0.0001). Mice fed the MHF diet with access to a running wheel weighed significantly less than the MHF-fed mice without a running wheel (WT-MHF-RW = 32.9 g ± 3.0) but they still weighed significantly more than the CF-fed treatment groups (Figure 1, A and C). The bodyweight of KO mice was not affected by wheel access or diet modification (KO-CF-SED = 25.4 g ± 1.3, KO-CF-RW = 26.1 ± 1.5, KO-MHF-SED = 25.6 ± 2.2, KO-MHF-RW = 25.2 ± 1.4) (Figure 1, B and C). The same pattern was present in the adiposity of the mice. WT mice fed the CF diet displayed similar amounts of adipose tissue regardless of wheel access (WT-CF-SED = 0.97 g ± 0.37; WT-CF-RW = 1.04 g ± 0.35). And the MHF-fed mice without a wheel displayed the most adiposity while the MHF-fed mice with wheel access were intermediate, with significantly more adipose tissue than the CF-fed groups and significantly less adipose tissue than the MHF sedentary mice (WT-MHF-SED = 6.44 g ± 1.39; WT-MHF-RW = 2.97 ± 0.68; F(7,49) = 85.43; p < 0.001) (Figure 1D). KO mice displayed similar amounts adipose tissue regardless of wheel access or diet modification (KO-CF-SED = 0.61 g ± 0.18; KO-CF-RW = 0.74 ± 0.38, KO-MHF-SED = 0.70 ± 0.23, KO-MHF-RW = 0.52 ± 0.21) (Figure 1D). KO mice displayed the slim body type and reduced adiposity reported in prior experiments, but this effect did not reach statistical significance in our cohort. This pattern is replicated again in the results of the IPGTT. Again, WT mice maintained on the CF diet with or without wheel access showed no differences in their ability to clear glucose, as assessed by calculating the area under the curve of their respective glucose tolerance curves (WT-CF-SED = 10,589 ± 4,196; WT-CF-RW = 9,073 ± 3,624). Significant glucose intolerance was present in the MHF-fed mice without wheel access while the MHF-fed mice with a wheel were again at an intermediate value between the CF treatment groups and the MHF sedentary group (WT-MHF-SED = 20,922 ± 5313; WT-MHF-RW = 15,462 ± 5572; F(7,76) = 20.27; p < 0.0001). All KO treatment groups displayed similar glucose clearance rates regardless of wheel access or diet, exhibiting the increased glucose sensitivity compared to WT previously reported, but again not reaching statistical significance in this metric (KO-CF-SED = 5,839 ± 2,945; KO-CF-RW = 6,348 ± 1,865, KO-MHF-SED = 5,932 ± 923, KO-MHF-RW = 6,367 ± 1,474).

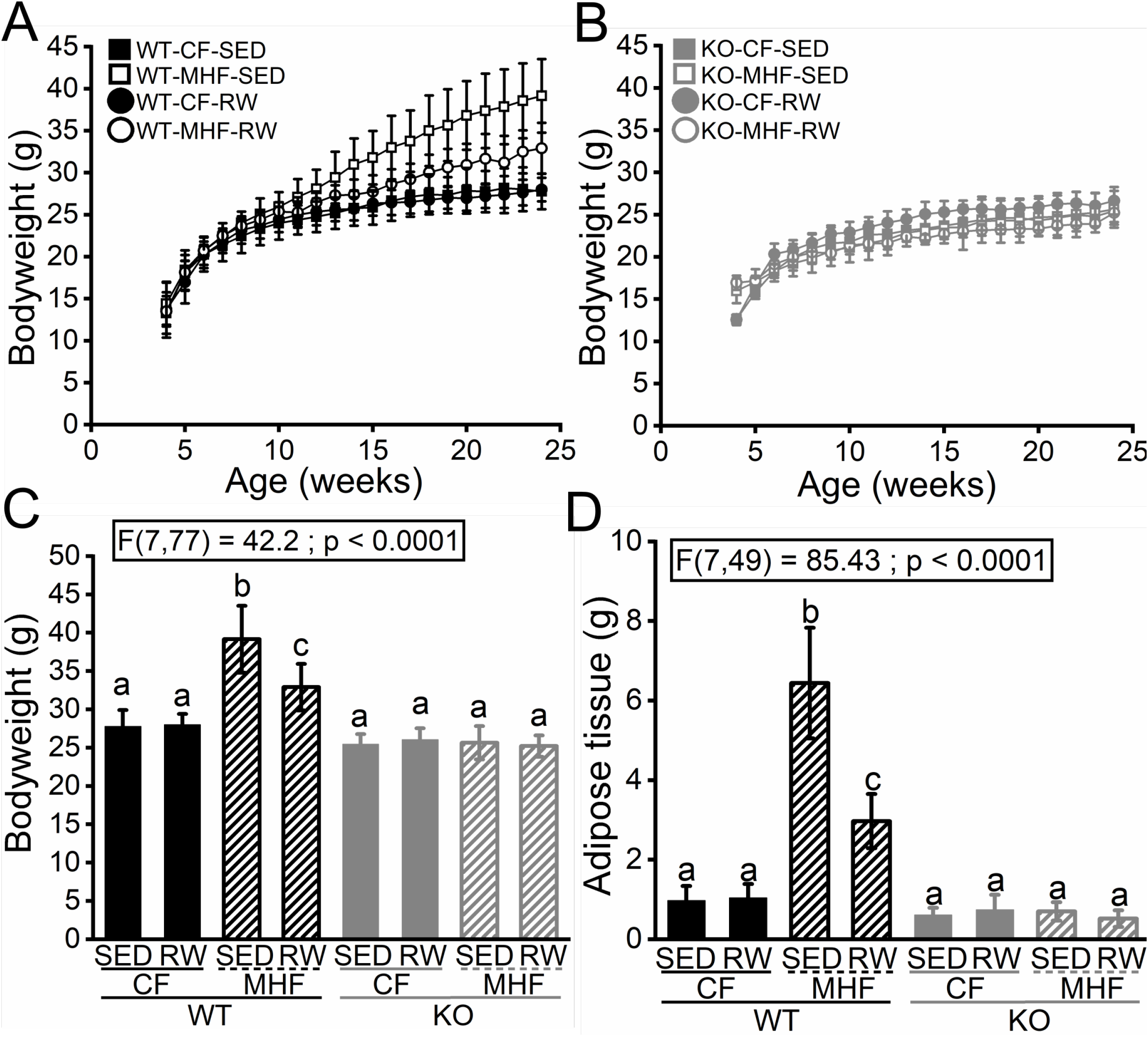

### Wheel running is consistent over time independent of genotype or diet

Following the 3-day acclimation period and barring any technical errors, mice in all treatment groups that were provided a wheel ran very consistently over the full 120-160 day running period. Qualitative assessment of double-plotted actograms showed that mice participate in wheel running in a manner that is entrained to their light/dark cycle with very little deviation (Figure 3). Tracking the mean daily running distance over time showed that on a weekly and monthly basis, the mice ran consistently (Figure 4C-D).

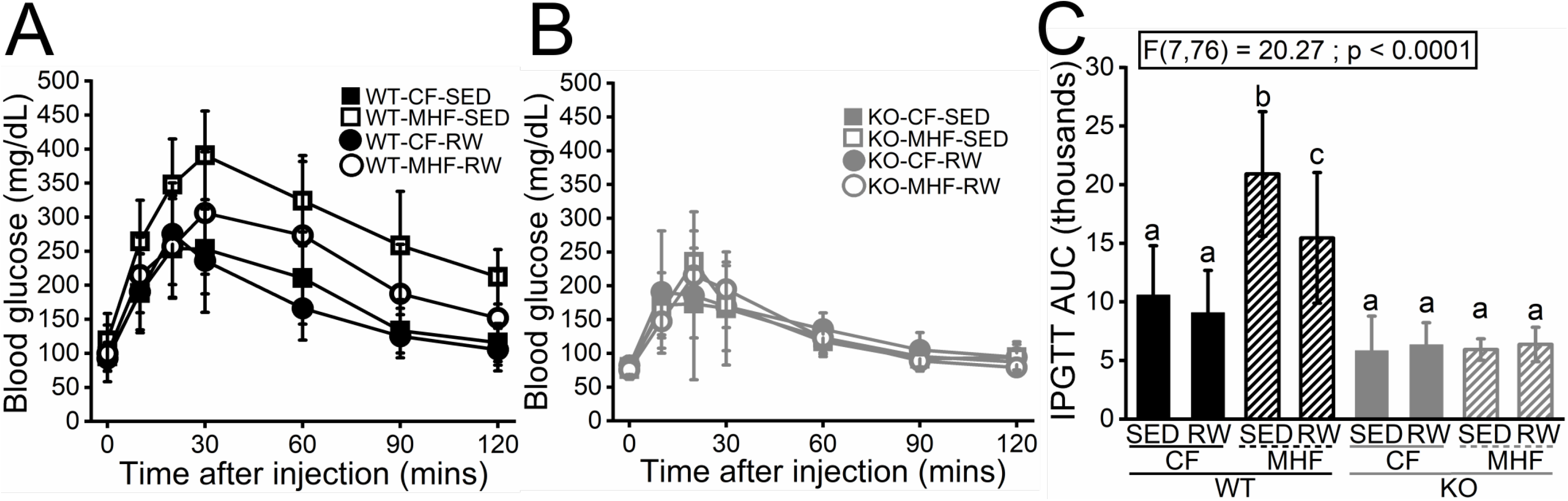

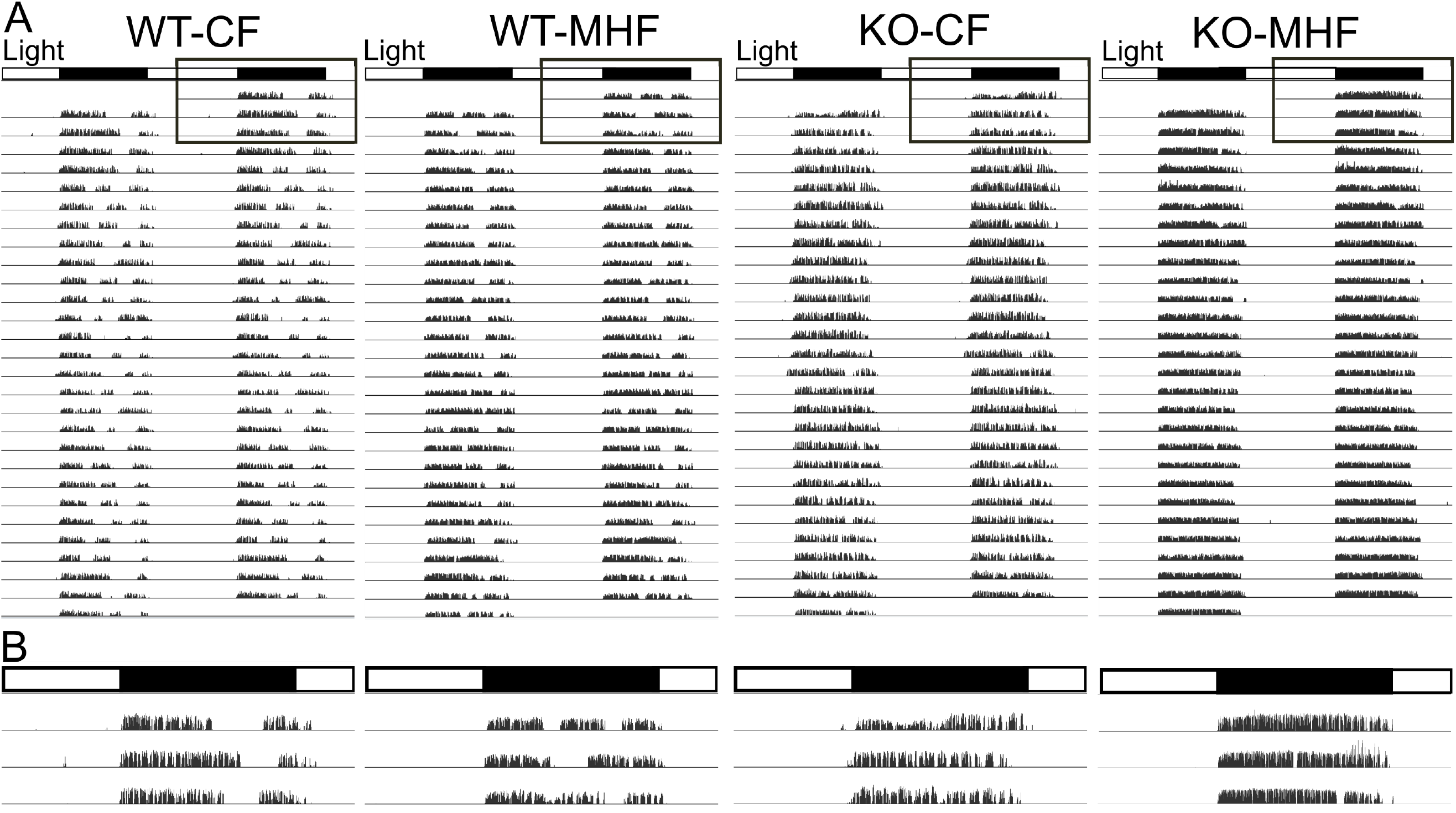

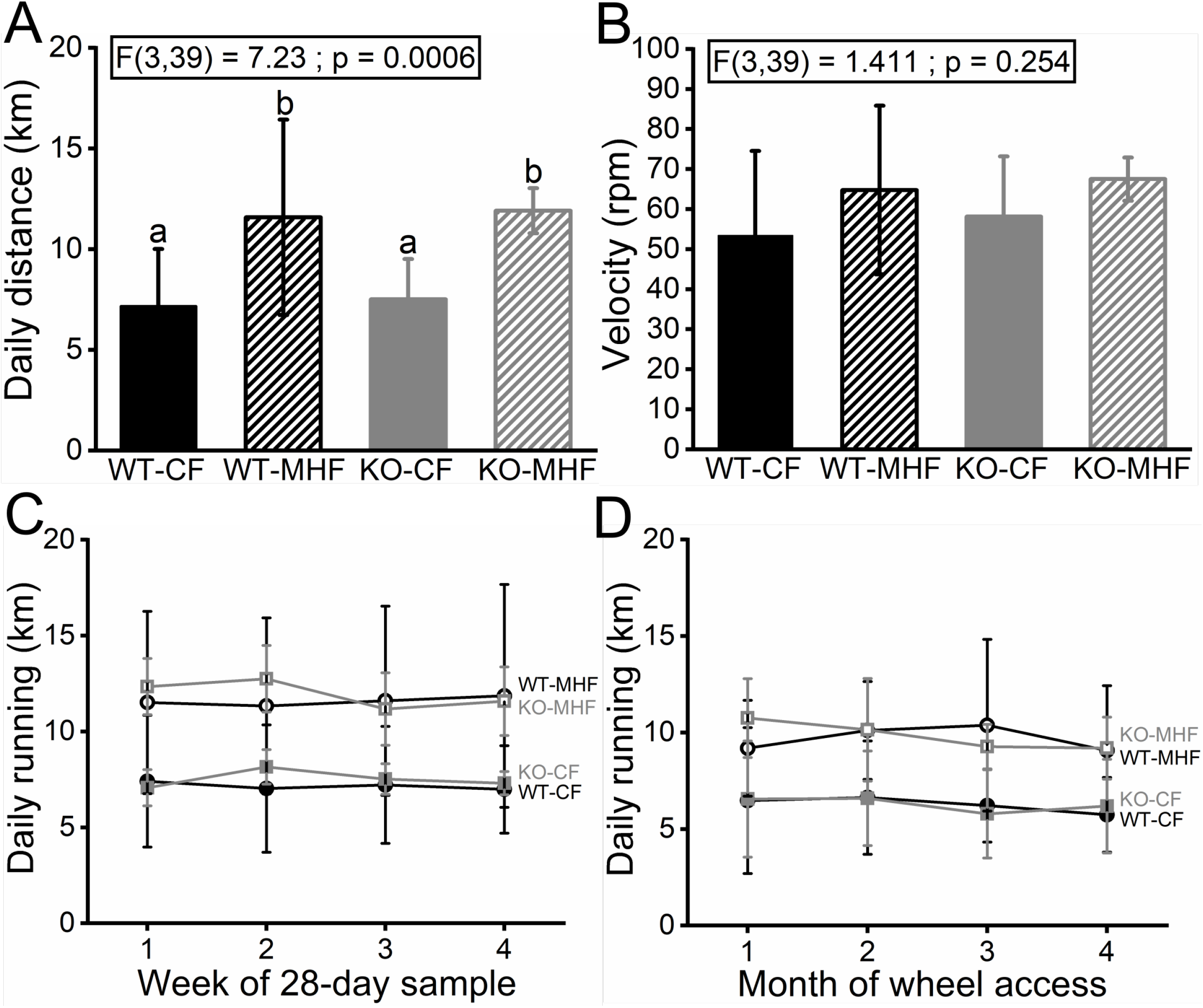

### K_v_1.3 ablation and fatty diet affect wheel running activity

Both WT and KO mice maintained on the MHF diet ran farther than the two CF-fed groups (WT-CF = 7.16 ± 2.85 km; KO-CF = 7.51 ± 2.0; WT-MHF = 11.58 ± 4.85; KO-MHF = 11. 91 ± 1.12; F(3,39) = 7.23, p = 0.0006) (Figure 4A). The WT-MHF and KO-MHF mice ran slightly faster than the WT-CF and WT-MHF but this did not reach statistical significance (Figure 4B).

We measured both the latency to start wheel running after the onset of the dark phase and the latency to stop wheel running after the onset of the light phase. All treatment groups began running soon after the onset of the dark phase (Figure 5C). WT mice continued running after the onset of the light phase whereas KO mice stopped running even before light onset (WT-CF = 13.7 ± 27.7 active minutes after light onset, WT-MHF = 25.6 ± 18.1, KO-CF = −52.3 ± 44.8, KO-MHF = −33.6 ± 12.2; F(3,39) = 17.2, P < 0.0001) (Figure 5D).

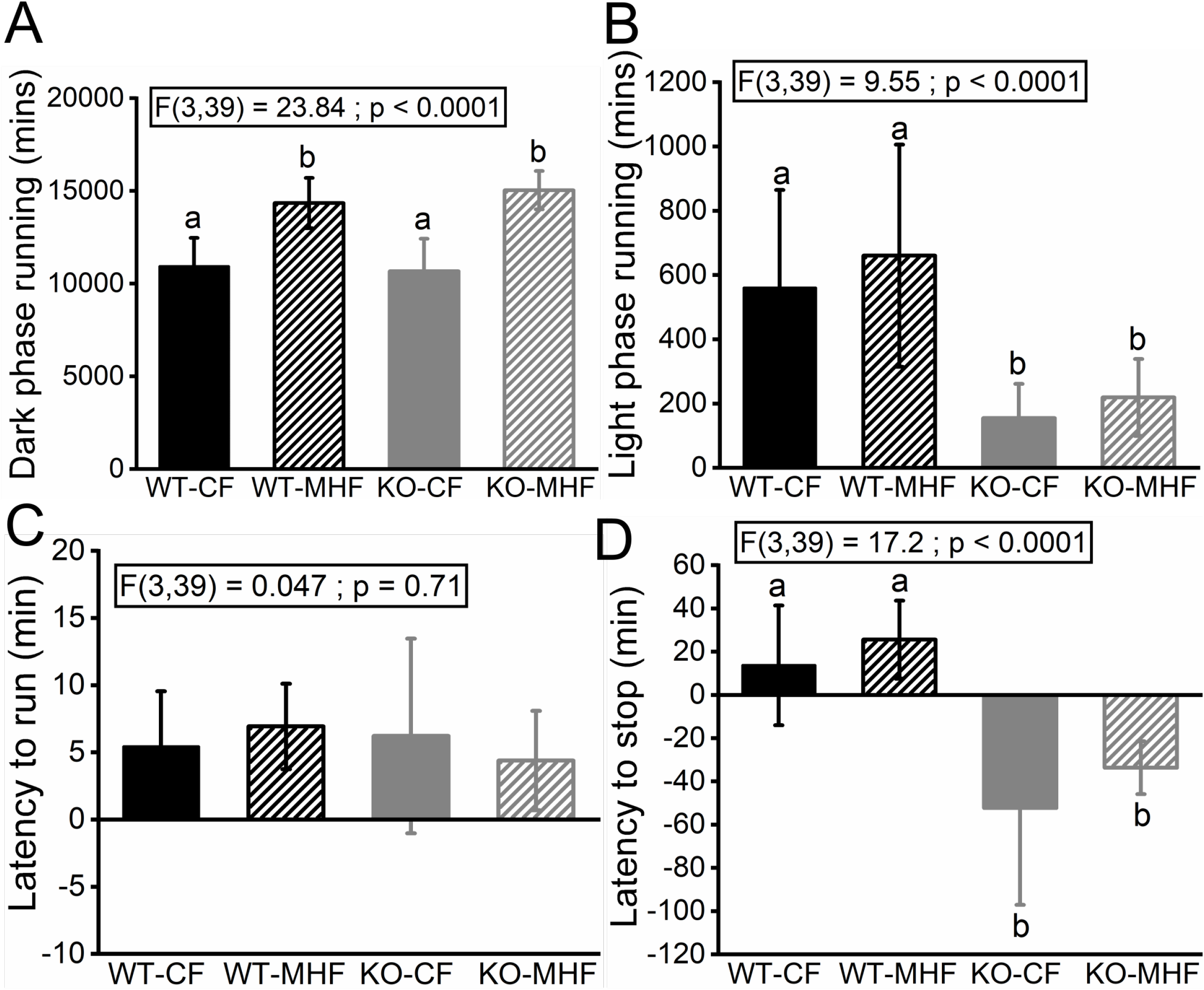

Across the 28-day sample of wheel running data, there are a total of 40,320 minutes, split evenly between the light and dark phases. We tracked how many minutes the mice were actively participating in wheel running in the light and dark phases. The MHF-fed mice spend more time running than the CF-fed mice across both genotypes (WT-CF = 10,927 ± 1,541 mins, KO-CF = 10,674 ± 1,754, WT-MHF = 14,347 ± 1,358, KO-MHF = 15,039 ± 1,034; F(3,39) = 23.84, p < 0.0001) (Figure 5A). In the light phase, the WT mice spent more time running than the KO mice across both diet treatments (WT-CF = 560 ± 305, WT-MHF = 660 ± 346, KO-CF = 155 ± 107, KO-MHF = 219 ± 120; F(3,39) = 9.55, P < 0.0001) (Figure 5B).

Running bursts were calculated as the number of consecutive 1-minute bins that a mouse was actively engaged in wheel running. Rests were calculated similarly as the number of consecutive 1-minute bins during which the mouse was not actively running. Thus, the length of a burst or rest could range from 1 minute long to hours. All four treatment groups engaged in the same number of bursts (WT-CF-RW = 1227 ± 252; WT-MHF-RW = 1218 ± 93; KO-CF-RW = 1282 ± 194, KO-MHF-RW = 1,093 ± 150; F(3,39) = 1.48, p = 0.23), but the length of the running bursts were significantly longer for MHF-fed mice compared to CF-fed mice for both genotypes (WT-CF-RW = 8.6 ± 1.6 mins per burst, KO-CF-RW = 7.6 ± 0.8, WT-MHF-RW = 11.5 ± 2.0, KO-MHF-RW = 13.5 ± 2.3; F(3,39) = 23.2, p < 0.0001)(Figure 6A+B). KO-MHF-RW mice engaged in the fewest rests, significantly less than the CF-fed groups but not statistically significant compared to the WT-MHF-RW group (WT-CF-RW = 1,217 ± 208, WT-MHF-RW = 1,156 ± 87, KO-CF-RW = 1,315 ± KO-MHF-RW = 1,005 ± 134; F(3,39) = 6.1, p = 0.002)(Figure 6C). Both MHF-fed groups engaged in shorter rests compared to the two CF-fed groups (WT-CF-RW = 7.9 ± 2.0 mins per rest; KO-CF-RW = 7.4 ± 2.0; WT-MHF-RW = 5.0 ± 1.0, KO-MHF-RW = 5.1 ± 0.9; F(3,39) = 8.8, p < 0.0001)(Figure 6D). We examined the distribution of resting behavior across the dark phase by binning the activity into hours. Independent of diet, both WT groups had a peak resting of 40 minutes 10 hours into the dark phase, which shortened toward the end of the dark phase, with concomitant increased activity. The KO mice never exhibited such a plateau and steadily increased rests periods incrementally throughout the dark phase up through the transition to the light phase (Figure 6E+F).

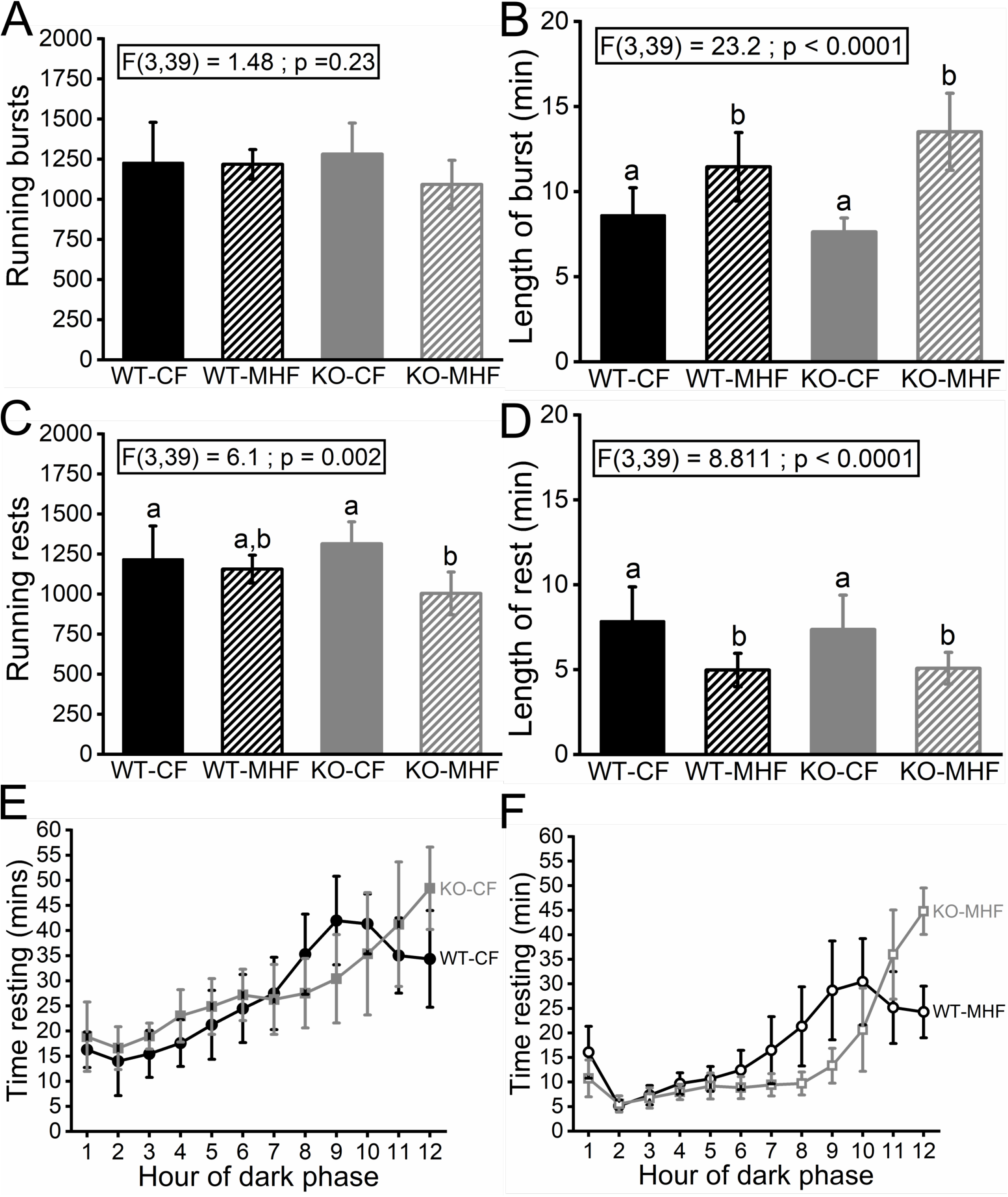

## DISCUSSION

These studies sought to investigate the effects of diet modification and genetic deletion of the voltage-gated potassium channel K_v_1.3 on voluntary exercise. K_v_1.3-null mice ran the same distance and speed as WT mice when they were maintained on the same diet. However, the patterns of running activity and resting were different in the two genotypes. Both genotypes were highly active on their wheels in the early portion of the dark phase, spending little time resting. Both genotypes also steadily increased the amount of resting as the dark phase progress. WT mice rested the most during hours 9 and 10, increasing their activity again in the last two hours of the dark phase while the KO mice did not exhibit this plateau, resting most heavily during hours 11 and 12 of the dark phase. In fact, the WT mice continued running even after the onset of the light phase while the KO mice seemingly predicted the transition to the light phase, ceasing their running behavior even before light onset. Previous studies have demonstrated increased energy expenditure and locomotor activity KO mice, but this is a novel characterization of their voluntary exercise phenotype {23}.

We also observed alterations of wheel running behavior following diet modification. Both WT and KO mice ran farther distances daily when maintained on the MHF diet compared to the CF diet. We determined that, in both genotypes, the increased daily running distance is primarily driven by MHF-fed mice participating in longer bursts of running and shorter rests compared to CF-fed mice. The MHF-fed mice did run slightly faster than their CF-fed counterparts, but this was not statistically significant. This is interesting because the body of literature is fairly split on this topic. Vellers et al. 2017 observed reduced time spent running, running distance, and running velocity in mice fed a high-fat, high-sugar diet in both short- and long-term feeding {37}. It is worth noting that in this experiment short-term is defined as a 3-day period of access to the high-fat, high-sugar diet and the long-term exposure was 14 days. Other studies have demonstrated similar reductions in mouse wheel running following maintenance on various high-fat diets while both mouse and human studies have demonstrated that increased caloric intake reduces spontaneous physical activity {38,39,40}. However, some investigators have observed an increase in wheel running following exposure to high-fat diet. For example, Meek et al. 2010, noted an increase in wheel running in a line of mice selective bred for high levels of voluntary exercise following a switch to a “Western diet.” {41}. Further, some studies have noted genotype-specific alterations of voluntary exercise, noting changes in some lines but not others following diet modification {42}. The contrasting effects observed between previous studies and the results we report here may be due to the amount of time the mice are maintained on the modified diet, the composition of the diet, or slight differences between the lines of mice being studied.

Participation in voluntary exercise partially ameliorated the negative metabolic effects of consumption of the MHF diet. WT mice with running wheel access maintained on the MHF diet showed lower bodyweight, lower adiposity, and improved glucose tolerance compared to WT mice on the same diet without a wheel. Our results replicated what has been demonstrated by previous studies investigating combinatorial effects of diet and exercise modification. Mice with access to a running wheel consistently display better metabolic health when maintained on MHF diet, but the benefits of the wheel are not as effective as removal of the MHF diet (or of never being exposed to the MHF diet to begin with). Thus, the high-fat fed mice given wheel access typically display an intermediate level of metabolic health that is between MHF non-runners and CF fed mice. Hicks et al. 2016 found lower weight gain and lower levels of glucose intolerance in mice fed a 60% kcal from fat diet after 11 weeks in the group that had running wheel access compared to the group that only had access to a locked wheel {43}. Yoshimura et al. 2018 observed that maintenance on a 57% kcal from fat diet induced accumulation of liver fat that was reduced following 1 week of wheel running and other studies have displayed similar results of at least partial rescues of varying negative metabolic phenotypes {44}. The KO mice are already resistant to weight gain, adipose deposition, and glucose intolerance and we did not observe any differences in these metabolic parameters regardless of diet or wheel access in the KO mice.

Our study has revealed that KO mice exhibit a voluntary exercise pattern that is different than WT mice in regards to their patterns of rest, especially later in the dark phase. We have also demonstrated that both WT and KO mice, when maintained on an MHF diet for 6 months, increase their levels of voluntary exercise compared to mice maintained on the CF diet. Understanding how these two variables affect exercise behaviors is important because of the growing potential of K_v_1.3 as a therapeutic target. We believe it is especially interesting because of the role that K_v_1.3 plays in the inflammatory response and the recent hypothesis that obesity is a state of chronic low-grade inflammation {45}. This means that K_v_1.3-targeted therapies, exercise regimens, and diet modification could be used in conjunction to treat metabolic disorders, so it is important to understand how these treatments interact.

Finally, our results suggest that it might be important for researchers using transgenic rodent models to monitor the running behavior of their animals in finer detail than is typically reported. The amount of exercise that mice perform each day is highly variable depending on the transgenic line being used {33}. Further, some studies do not even report the amount of exercise and seem to assume that all mice are running normally simply because a wheel is present in their cage. If investigators only quantify daily activity (or don’t quantify activity at all), they could be missing differences in exercise patterns that are potentially physiologically relevant.

## Author Contributions

B. Chelette and D. Fadool conceived of the study, analyzed the data, and wrote the manuscript. B. Chelette and A. Thomas performed experiments, collected system physiology measurements, and monitored voluntary exercise. All authors performed animal husbandry and contributed to the final version of the manuscript.

## ACKNOWLEDGEMENTS

We would like to thank Daniel Gonzalez, Margaret Vinson, and Kassandra Ferguson for their technical assistance in maintaining running wheels, measuring body weights and collection of adipose tissues. We would like to thank Carley Huffstetler for routine laboratory management and mouse husbandry.

